# Overlap in the cortical representation of hand and forearm muscles as assessed by navigated TMS

**DOI:** 10.1101/2023.01.27.525857

**Authors:** Fang Jin, Sjoerd M. Bruijn, Andreas Daffertshofer

## Abstract

The representation of upper limb muscles in the motor cortex is complex. It contains areas of excitability that may overlap between muscles. We expected the cortical representations of synergistic muscle pairs to overlap more than those of non-synergistic muscles. To detail this, we used navigated transcranial magnetic stimulation in eight hand and forearm muscles of twenty healthy participants. We transformed the cortical representations of muscles to a template MRI to allow for group analysis. We found that the amount of overlap in cortical representations differed significantly between within-hand and within-forearm muscle combinations. Most synergistic muscle pairs, both within the hand, within the forearm and between them, had a larger overlap than non-synergistic muscle pairs. Our study supports the largely overlapping nature of cortical representations of upper limb muscles. We can particularly underscore that the overlap is elevated in muscles that usually act in a synergistic manner.

## Introduction

Penfield and Boldrey [1] localised the somatotopic representation of motor processing in the primary cortex (M1) using electrical stimulation. While many subsequent studies established the principle structure of this motor homunculus, more fine-grained approaches revealed that it might deviate from a well-ordered organisation in that especially upper limb muscles can have complex and sometimes overlapping representations [2]. Shinoda, Yokota, and Futami [3] demonstrated axon collaterals from a single corticospinal neuron to branch into motor neuron pools of at least four muscles. More recent imaging studies confirmed that cortical representations of distinct upper limb muscles may overlap using either functional magnetic resonance imaging (fMRI) [4, 5] or transcranial magnetic stimulation (TMS) [6-9]. The overlap in the cortical representation of muscles may indicate synergistic control of muscle groups [10]. Its functional relevance has been underscored by Tyč and Boyadjian [11], who found an increase in the overlap between distal and proximal upper limb muscles due to motor training. Interestingly, different pathologies may come with an increased overlap of cortical representations, not necessarily restricted to the upper extremities [12-15]. According to Yao and co- workers [12], for instance, stroke may be accompanied by an overlap between elbow and shoulder cortical representations associated with the loss of independent control for elbow and shoulder, a hallmark for compensatory strategies in stroke survivors. Elgueta-Cancino et al. [16] reported the overlap between deep and superficial fibres of the multifidus muscle to be correlated with the severity of low back pain.

As a non-invasive technique, TMS is well suited to investigate the overlap in cortical representations of muscles. For instance, Melgari et al. [17] employed TMS to assess the amount of overlap between twelve muscles. They found a more pronounced overlap for hand-hand and forearm-forearm muscles combinations than between hand and forearm muscles, let alone for upper arm muscles. More recently, Tardelli et al. [18] reported representations of forearm muscles to overlap more than those of intrinsic hand muscles. Muscles that are known to be active in unison seem to have more cortical overlap than others [18]. Especially synergistic muscles seem to overlap more than non-synergistic ones [10].

To further unravel the signature of synergistic muscle combinations, we investigated the cortical representations of eight hand and forearm muscles in twenty healthy volunteers using navigated single-pulse TMS eliciting motor-evoked potentials (MEPs). To warrant a sufficient amount of MEPs as well as proper spatial resolution, we employed a pseudo-random TMS positioning [19]. We analysed the areas of excitability per muscle on subject-specific cortical surfaces of up to 1 mm resolution [20, 21]. In [20] we showed that both the size of the surface area and its centroid are valid and reliable metrics. However, we also realised that both come with considerable between-subject variability arguably because of clear differences in individual brain anatomies. This variability may jeopardise group analysis, for which we standardise anatomy by warping the subject-specific surfaces as well as the points of TMS focus to the common MNI template.

We hypothesised the muscles to have distinct, albeit partially overlapping cortical representations. We expected the overlaps between hand-hand, hand-forearm, and forearm-forearm muscle pairings to differ and, in particular, synergistic muscles to show a larger overlap than non-synergistic ones.

## Methods

Twenty healthy adults (eight females) participated in the study. Before experimental assessments they filled out standard TMS and MRI screening questionnaires and were informed of the measurement procedures and risks. All participants provided signed informed consent. The study had been approved by VUmc Medical Ethics Committee (2018.213 - NL65023.029.18). The experiment was conducted in line with the Declaration of Helsinki.

### TMS measurement

Prior to TMS application, all the participants underwent T1-weighted MRI scanning (3T Achieva, Philips, Best, The Netherlands; matrix size 256×256×211, voxel size 1.0×1.0×1.0 mm^3^, and TR/TE 6.40/2.94 ms). We integrated the anatomical scans in the neuro-navigation system (Neural Navigator, Brain Science Tools BV, De Bilt, The Netherlands, www.brainsciencetools.com) by segmenting them for grey matter using SPM (SPM12, https://www.fil.ion.ucl.ac.uk/spm/software/spm12/) and identifying four fiducial points (nasion, nose tip, left and right peri-auricular points) for co-registration.

Single-pulse mono-phasic stimulations were delivered using a Magstim 200^2^ TMS stimulator with a 70 mm diameter figure-of-eight coil (Magstim Company Ltd., Whitland, Dyfed, UK). The elicited MEPs were captured by a 16-channel EMG amplifier (Porti, TMSi, Oldenzaal, the Netherlands) and sampled at 2 kHz. Bipolar electrodes were positioned following SENIAM convention; see Figure 1.

**Figure 1.**
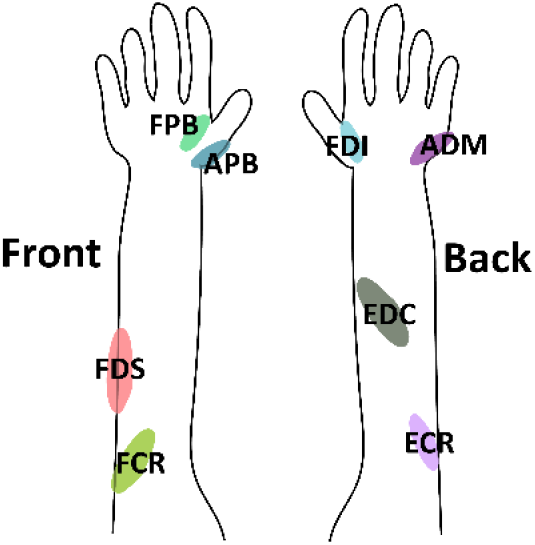
Eight muscles of the right hand and right forearm considered in this study: first dorsal interosseous (FDI), abductor digiti minimi (ADM), flexor pollicis brevis (FPB), abductor pollicis brevis (APB), extensor digitorum communis (EDC), flexor digitorum superficialis (FDS), extensor carpi radialis (ECR), and flexor carpi radialis (FCR). In the left panel the palm is pointing inward, in the right panel it faces outward.

After identifying the respective hot spots, we determined the resting motor thresholds (RMTs) for FDI, EDC, and FCR, following our procedures outlined in [20]. In brief, for each of the three muscles we first stimulated with increasing intensity until a consistent MEP appeared. Then, we applied thirty stimulations at around central gyrus (M1) from, which the points with the highest MEP amplitude was considered as hot spot. At the hot spot, the RMT was determined as the lowest intensity at which the MEP amplitude exceeded 50 μV in five out of ten stimulations. The three RMTs served to define three conditions, namely stimulation intensities set to 105% of the respective RMTs. We deliberately chose such low stimulation intensities to limit the affected cortical region [21] and, by that, to avoid spurious overlaps between cortical representations at higher intensities [22].

Per intensity we stimulated 120 times with a 5s inter-stimulation duration (fixed via a revised version of https://github.com/armanabraham/Rapid2). This procedure we repeated once yielding at total of 3 (intensities) × 120 (stimulations) × 2 (repetitions) = 720 stimulations per participant. As already mentioned, we employed a pseudo-random coil positioning [19] that covered roughly 5×5 cm around the corresponding hot spot.

### Data analysis

The EMG signals were high-pass filtered at 30 Hz using a 2^nd^-order bi-directional Butterworth filter. We identified stimulations with proper motor-evoked potential (MEPs) based on the peak-to-peak EMG amplitude (less than 10 mV but larger than twenty times the baseline’s standard deviation obtained 200 ms prior to stimulation). All stimulations were classified as either MEP or non-MEP points. Stimulations outside M1 (“precentral L” in the “Mindboggle6” atlas [23]) were eliminated; see also the *Supplementary Section* S7 for the analysis without this constraint.

Our data analysis relied on the individual MRIs that were segmented using FreeSurfer (http://surfer.nmr.mgh.harvard.edu/) and we used the pial-surface for all subsequent steps. For the within group comparison we used the MNI152 default subject implemented in Brainstorm [24] as template surface; see *Supplementary Sections* S6 and S7 for the analyses on the subject-specific MRIs.

The subsequent analysis steps are illustrated in Figure 2. Coil coordinates and orientations were registered from the individual MRI surface (panel B) and we inflated each hemisphere to a unit sphere [25,26] (panel C). To map a stimulation to the template (panel D), we searched for the sphere-inflated template vertex by incorporating subject-specific gyri and sulci locations via their curvature values and warped to the respective inflated (spherical) template by minimising their great-circle distance. The resulting points (panel D) were finally deflated to a template surface (panel E).

**Figure 2.**
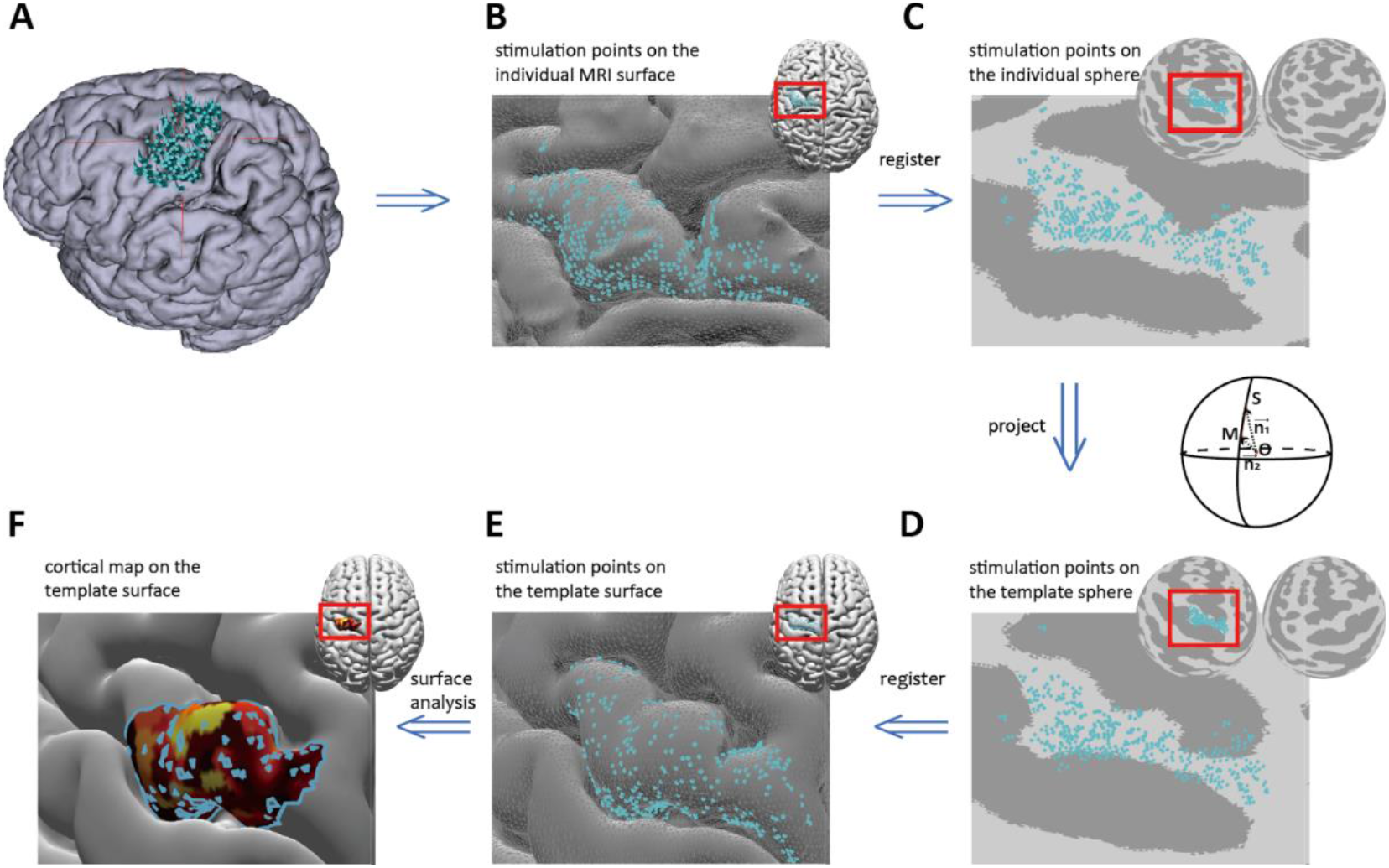
Processing pipeline. The blue dots represent the stimulation points (denote by *S* on the sphere). The surfaces in panels A and B are the individual MRI surfaces and panel C depict the mapping to the unit spheres per hemisphere. The lower row relates to the template representation where panel D is the transform to the corresponding unit spheres that can readily mapped to the MNI152 template (panel E). Panel F displays the estimated area of excitability, colour-coded by the MEP amplitudes.

We used our open-source surface analysis toolbox (https://github.com/marlow17/surfaceanalysis) [20] to quantify the cortical representation of every muscle by its area. In the triangulated cortex mesh, we first determined the area’s vertices, i.e., 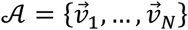 with 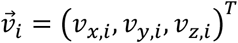 and set their MEP values MEP_*i*_ by interpolating the MEP amplitudes corresponding to the original stimulation points – see [20] for further details about the underlying search algorithm. Here, we abbreviate the resulting (MEP-weighted) area by 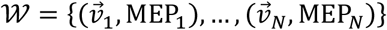. In, e.g., Figure 2 (panel F), the area is shown as colour-code patches with yellow indicating the largest MEP amplitudes, i.e., the highest degree of muscle-specific excitability.

Following [20], we first parametrised an area’s location via its centroid given by

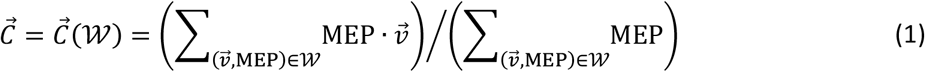

Next, we computed the size of an area via the triangular prism. For this, consider a triangle in *𝒜* let the lengths between its vertices 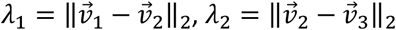 and 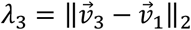, where ∥…∥_2_ denotes the Euclidean distance. Then, the area size *W :=* ∥*𝒲*∥ reads

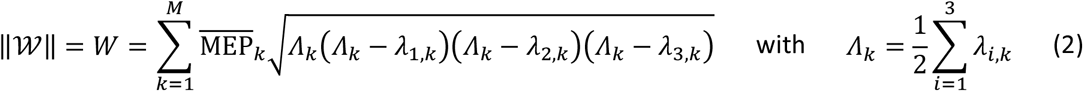

with 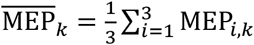 being the mean value of (interpolated) MEP amplitudes at the three vertices of triangle *k*.

Finally, we assessed the overlap of the cortical representations by combining the eight muscles into 28 distinct pairs (FDI-ADM, FDI-APB, …) and defining three groups of muscles: hand-hand, hand-forearm, and forearm-forearm. This grouping eased focusing on effects of (non-)synergistic muscle combinations. For instance, within the hand-hand group, FDI-APB, FDI-FPB, and APB-FPB are typically considered synergistic, and in the forearm-forearm group EDC-ECR and FDS-FCR are typically considered synergistic. In *Supplementary Table* S1.1 we provide a complete list of muscle pair that we here considered as synergistic muscle combination and motivate our choice by providing the corresponding references. Yet, one has to realise that these combination strictly speaking depend on the motor task being studied.

Evidently, the overlap between areas can be quantified via their intersect that we here normalised using the corresponding union. That is, per muscle pair (*k, l*) we defined :

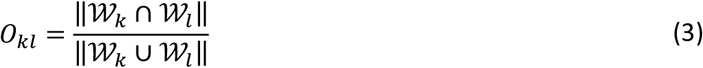

where ∥… ∥ denotes the size definition given in Eq. (2). This computation is illustrated in Figure 3. Here we would like to note that by transforming the subject-specific cortex stimulation points to the template (cf. Fig. 2) both 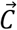 and *O*_*kl*_ could readily enter our group analysis (see below under *Statistics*).

**Figure 3.**
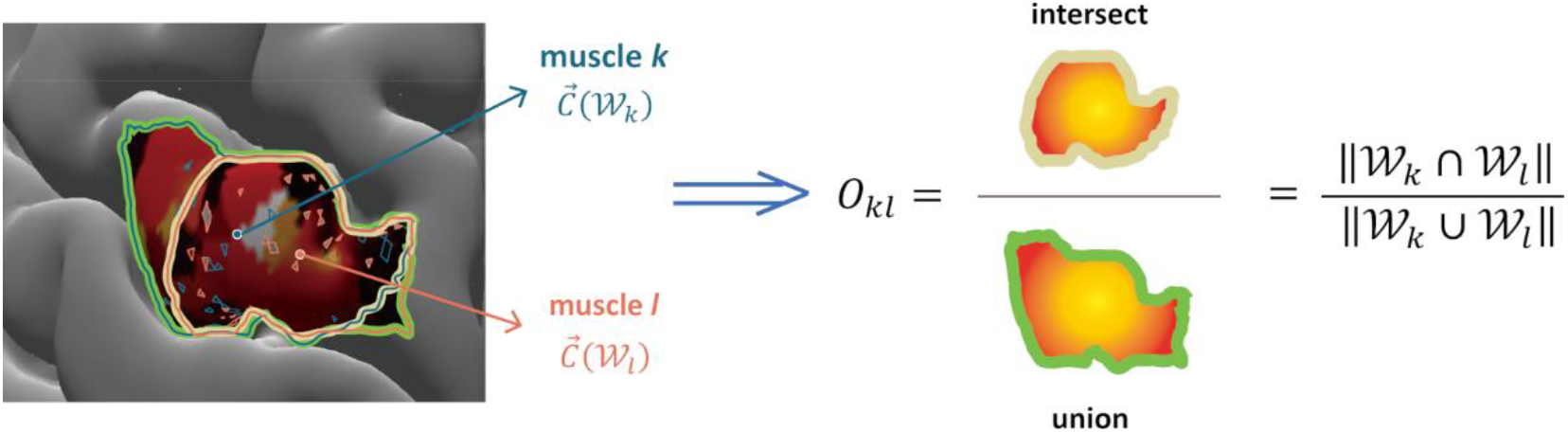
Illustration of the centroid and overlap definition of two muscles k and l. To ease visual inspection, we highlighted the contour of the two areas; cf. Eq. (3).

### Statistics

A two-way ANOVA with factors of *Muscle* and *Intensity* served to test for significant differences in the centroids 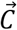 and the area size *W* across all muscles. Subsequently, we grouped the muscles into hand and foreman muscles and analysed between-group and within-group (hand-hand, forearm-forearm, and hand-forearm) differences in the overlaps *O*_*kl*_ using a two-way ANOVA with factors *Muscle-Group* and *Intensity*. For the overlaps, we also performed post-hoc tests of muscle combinations within groups of pairs. For this we conducted three separate two-way ANOVAs with *Pair* and *Intensity* as factors. Note that the results of the post-hoc analysis of the all combinations are reported in tabular form in *Supplementary Material* S5. Prior to analyses, we tested for sphericity using Mauchly tests and applied a Greenhouse-Geisser correction whenever necessary. Throughout analysis we considered a significant threshold of α = 0.05; all ANOVAs used repeated measures. Post-hoc assessments were Bonferroni corrected. We realised all analyses in Matlab 2022a (MathWorks, Natick, MA, USA).

Please note that the number of elicited MEPs has already been reported in [20]; see there *Supplementary Table* S1 and *Supplementary Figure* S1.

## Results

### Muscle representations

Before summarising the results of our hypothesis testing, we first illustrate two examples of warping the cortical surfaces to the MNI152 template surface in Figure 4.

**Figure 4.**
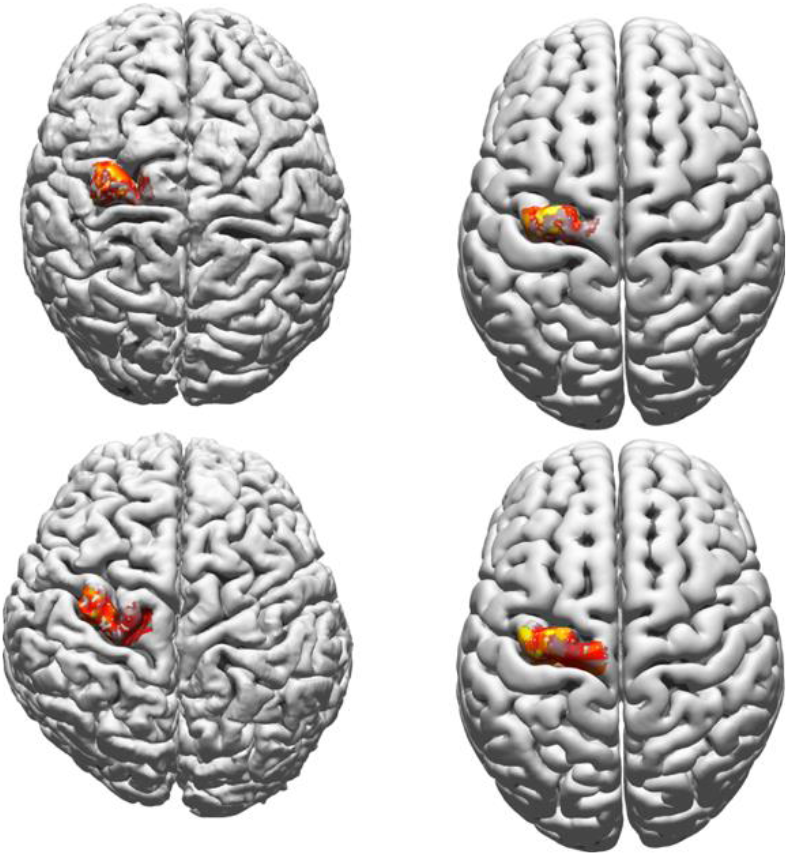
Two examples of the warping of the area of excitation of a single muscle to the template surface – left column: subject-specific cortical surface; right column: mapping on the template surface. One participant is of European descent (upper row), the other of East-Asian descent (lower row); in both cases the MEP-weighted region of excitability of FDI has been selected. Here we would like to note that stimulation points outside the contralateral M1, defined as the left precentral area, have been removed; see Fig. 2 for the corresponding procedures. Yellow indicates large MEP-weighted value, dark red low ones; units are arbitrary due to scaling to a maximum value of 1; cf. Figure 5.

Figure 5 depicts the corresponding group average for the 105%-RMT-FDI Intensity (= 47.25±2.16% stimulator intensity; EDC = 47.68±2.23% and FCR = 50.58±2.28% can be found in as *Supplementary Figures* S2.1 and S2.2). The representation clearly differed between muscles with, on average, FDI showing the highest degree of excitability.

**Figure 5.**
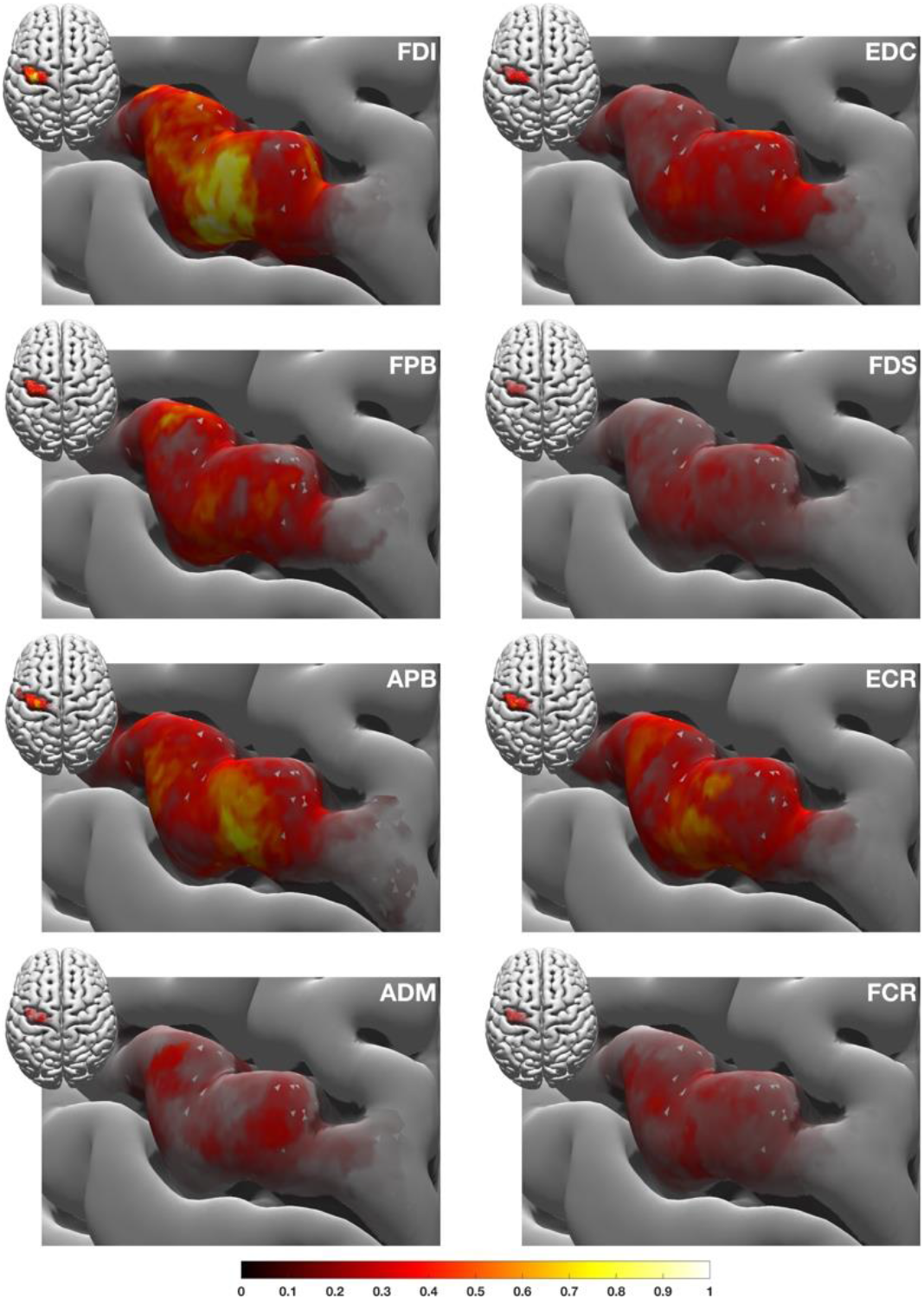
Group average of the cortical representations of the eight muscles under study. The areas are shown on the MNI template surface and represent excitability maps for the 105%-RMT-FDI intensity. The colour coding is given by the size of the corresponding MEPs, where yellow implies a large MEP amplitude and dark red a low one (the darker the colour the more transparent the plot will be to ease legibility); units are arbitrary thank to scaling. The same figure with muscle-specific scaling is given in the *Appendix* (Figure A1). In the *Supplementary Section* S2 we show the other two stimulation intensities as well as intensity maps of some randomly selected participants (*Supplementary Section* S3).

### Position and size of the cortical muscle representation

Overall, the hand representations were more lateral than those of the forearm muscles; see Figure 6 where we show the average centroid positions for the 105%-RMT-FDI in the 2D-plane (anterior/posterior vs. medial/ lateral direction).

**Figure 6.**
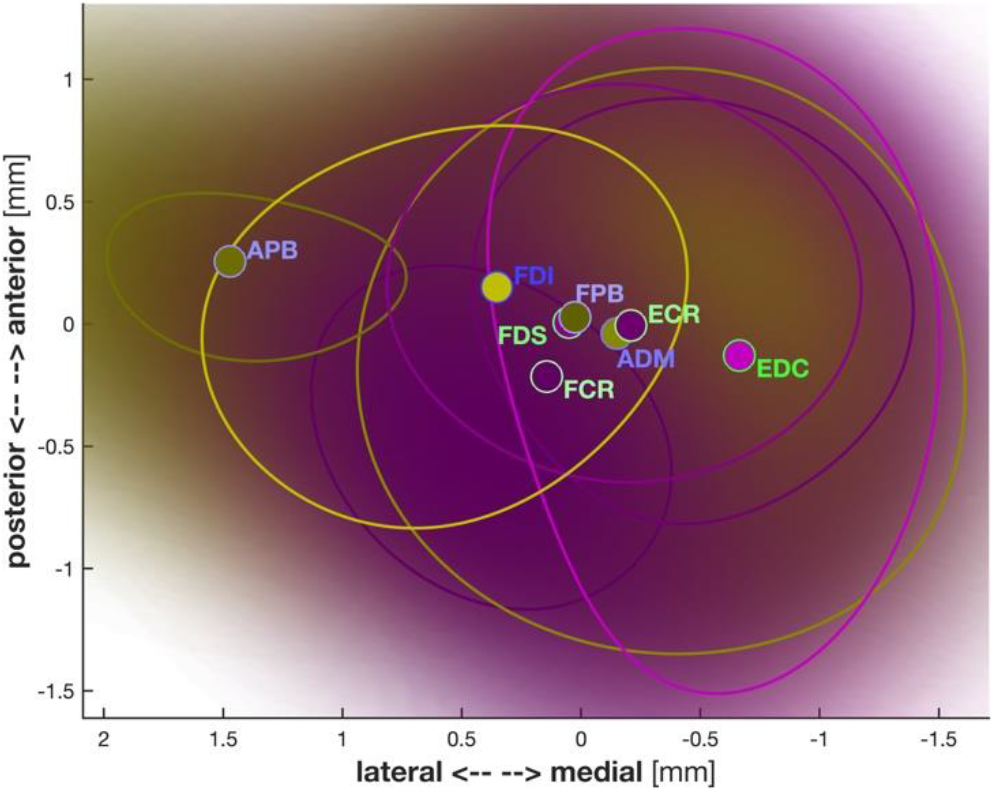
Probability densities of the centroids of the area of excitability estimated using kernel source densities over all participants. Contour lines indicate the 5% probability values. The different colours symbolize the hand (yellow) and forearm (magenta) muscles; dots are the mean centroids labelled accordingly. The hand muscles are more lateral than the forearm muscles. Note that we only depict the (anterior/posterior vs. medial/ lateral direction)-plane.

As summarised in Table 1, the centroid positions in the medial/lateral and inferior/superior directions (*C*_*y*_ and *C*_*z*_, respectively) as well as the area size differed significantly per *Muscle*. Neither parameter showed significant effects of *Intensity* or a significant *Intensity* × *Muscle* interaction.

**Table 1.** Statistics the cortical representation of the eight muscles. *W* = area size, centroid components: *C*_*x*_ = medial/lateral, *C*_*y*_ = anterior/posterior, *C*_*z*_ = inferior/superior. Significant effects are shown in **bold**.

| | *Intensity* | | *Muscle* | | *Intensity* × *Muscle* | |
| --- | --- | --- | --- | --- | --- | --- |
|  | F(2,26) | *p* | F(7,91) | *p* | F(14,182) | *p* |
| $C_x$ | 0.354 | .705 | 1.349 | .273 | 1.041 | .405 |
| $C_y$ | 0.090 | .914 | 4.668 | **.000** | 1.299 | .270 |
| $C_z$ | 0.760 | .435 | 4.728 | **.000** | 1.272 | .288 |
| $W$ | 3.865 | .060 | 3.147 | **.040** | 1.461 | .251 |

### Size of the overlapping areas

The overlap between representations differed significantly between muscle groups (F(2,38) = 208.862, *p* = .000). There was also a significant effect of *Intensity* (F(2,38) = 6.246, *p* = .005) as well as a significant *Intensity* × *Muscle-Group* interaction (F(4,76) = 5.326, *p* = .001). For the 105%-RMT-FCR Intensity, the overlap was larger than in both 105%-RMT-FDI (*p* = .021) and EDC (*p* = .011). Post-hoc pairwise comparisons revealed no significant differences in the overlap between the different pair groups. Corresponding descriptive statistics are illustrated in Figure 7.

**Figure 7.**
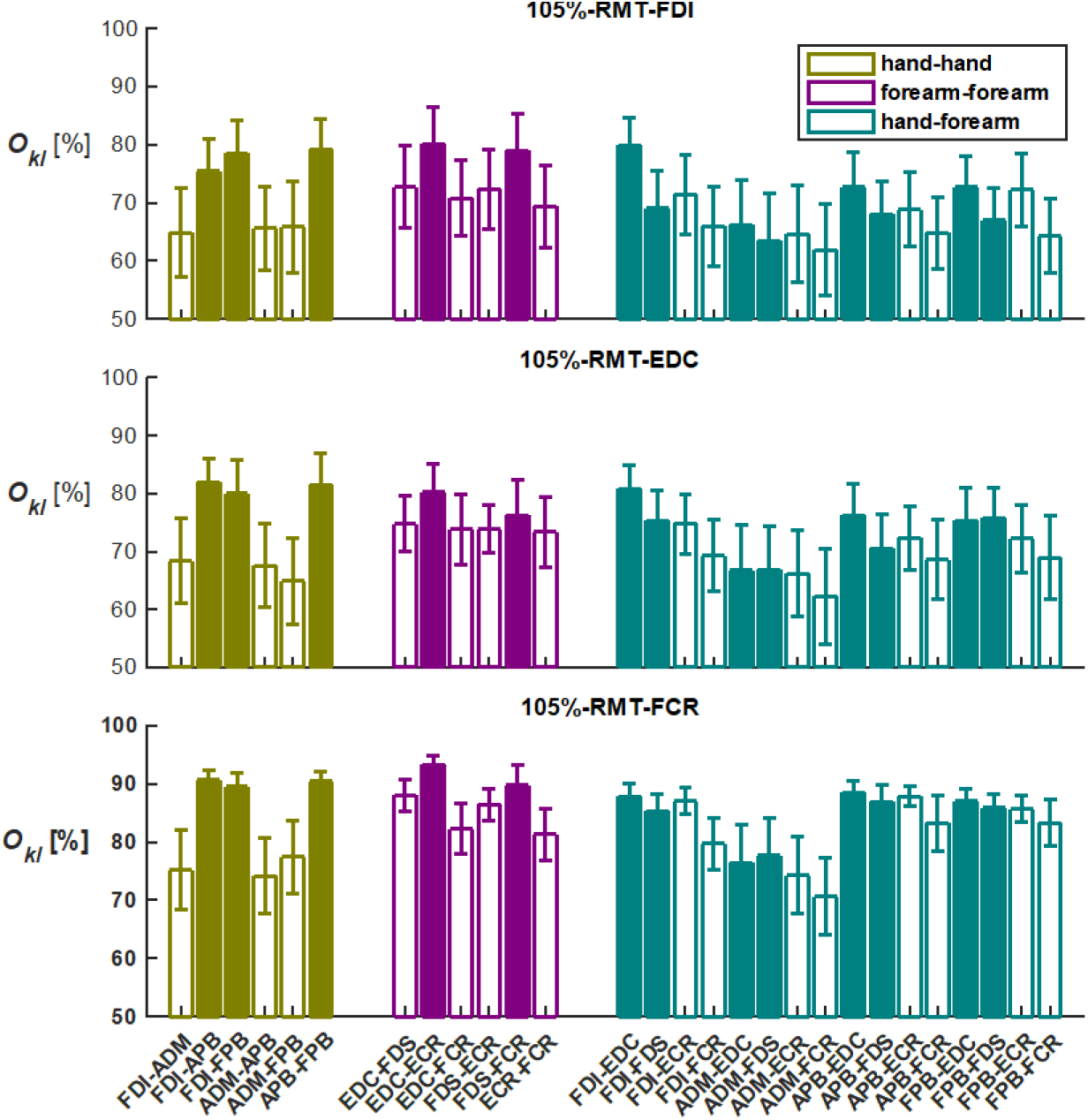
Overlap of all muscle pairs. The shown percentages of Overlaps O_*kl*_ between muscle pairs expressed in percentage (of the union area) for the three intensities. The filled bars represent the muscle pairs with synergistic function, whereas the open bars represent the non-synergistic muscle pairs; see Supplementary Table S1.1 for a motivated definition of synergistic pairs. Error bars represent the standard error over participants. Colour coding agrees with Figure 5.

### Within muscle-group overlaps (post-hoc analyses)

For the hand-hand muscle pairs the overlap displayed significant main effects of *Intensity* and *Pair* (F(2,38) = 4.979, *p* = .012 and F(5,95) = 4.021, *p* = .034, respectively) while the *Intensity* × *Pair* interaction did not reach significance (F(10,190) = 0.422, *p* = .819). The overlaps at 105%-RMT-FCR were larger than at 105%-RMT-EDC (*p* = .044). Analysing the forearm-forearm muscle group revealed pretty much the same effects (*Intensity*: F(2,38) = 3.768, *p* = .032 and *Pair*: F(5,95) = 3.870, *p* = .016 as well as *intensity* × *Pair*: F(10,190) = 0.433, *p* = .808). Again, the overlaps were larger for 105%-RMT-FCR than for 105%-RMT-EDC (*p* = .029). Additionally, the overlap of the FDS-FCR combination exceeded that of ECR-FCR (*p* = .027). Finally, the overlaps of hand-forearm muscle pairs were significantly affected by the factor *Intensity* (F(2,38) = 6.987, *p* = .003) but not by *Pair* (F(15,285) = 1.859, *p* = .164). The *intensity* × *Pair* interaction was again not significant (F(30,570) = 0.668, *p* = .706). However, here both 105%-RMT-FDI and -EDC intensities showed smaller overlaps than 105%-RMT-FCR (*p* = .012 and *p* = .013, respectively).

## Discussion

We mapped eight hand and forearm muscles using navigated TMS and assessed the overlap in cortical representations of various muscle pairs. Using surface analysis after warping the subject-specific cortex surfaces to a standardised template, we quantified the areas of excitability by centroid positions and area size. The hand representations were lateral, whereas those of the forearm muscles were more medial. This agrees with earlier studies that showed digit movements to be more laterally represented than upper limb movements [27]. We can also support that the thumb muscles (FDI, APB and FPB) are lateral to the little finger muscle (ADM).

### Overlap in cortical representations – synergistic versus non-synergistic muscles

Previous research using electrical stimulation in monkeys and humans demonstrated that cortical representations of the upper limb muscles can overlap [2]. As such we expected the cortical representations to be (partly) overlapping. Yet, we hypothesised the overlap to differ between the hand-hand, forearm-forearm, and hand-forearm muscle combinations as well as between synergistic and non-synergistic muscles. As summarised in *Supplementary Table* S1.1., we considered FDI-APB, FDI-FPB, and APB-FPB as synergistic piar within the hand-hand group. This especially applies for pick-up actions. The forearm-forearm pairs EDC-ECR and FDS-FCR are usually synergistic for larger extension and flexion, and in the hand-forearm group, EDC-FDI, -APB, -FPB, -ADM, and FDS-FDI, -APB, -FPB, -ADM usually act in a synergistic manner. Having said that, one must generally realise that the definition of synergistic muscle combination is typically task specific.

The amount of overlap did indeed differ significantly between hand-hand, hand-forearm, and forearm-forearm muscle pairs groups. Melgari et al. [17] showed that the amount of overlap in the hand-hand, hand-forearm, and forearm-forearm pairs was higher between hand-arm and arm-arm pairs. And according to Tardelli et al. [18], the overlap between intrinsic hand muscles (ADM-FPB) is smaller than between forearm and hand muscles (FCR-ADM and FCR-FPB). Our research revealed a significant difference between muscle-pair groups but not always so in pair-wise comparisons. This discrepancy calls for future studies involving even more muscles than we currently did, as this may clarify further how the amount of cortical overlap relates to the muscles’ anatomical locations.

By and large, the synergistic muscle pairs overlapped more than the muscle pairs without synergistic function. In the hand-hand muscle pairs, the average overlap of synergistic muscle pairs (FDI-APB, FDI-FPB, APB-FPB) was higher than the overlap in non-synergistic muscle pairs (ADM-FDI, ADM-APB, ADM-FPB). The synergistic muscle pairs (EDC-ECR, FDS-FCR) demonstrated more overlap than other forearm-forearm muscle pairs. The pair-wise comparisons revealed that FDS-FCR overlapped significantly more than ECR-FCR. Massé-Alarie et al. [10] also reported the ECR-EDC overlap to be stronger than the ECR-FCR overlap. Likewise, DeJong et al. [22] showed that the synergistic pairs FDI-APB and EDC-ECR overlap more than other muscle pairs.

The ADM-related muscles pairs (ADM-FDI, ADM-APB, and ADM-FPB) had smaller overlaps than the FDI-APB, FDI-FPB, and APB-FPB pairs. The average centroid of ADM was more medial than the centroids of FDI, APB, and FPB; see Figure 6. However, there was no significant difference in the centroids of cortical representation between other muscle pairs (ECR-FCR, ADM-FCR, etc.). We hence speculate that the observed overlaps between hand muscles may be associated with the distribution of the individual cortical representation. By contrast, the overlaps between forearm muscles seem to resemble (the structure of) other elements of the descending motor pathway.

### Standarising anatomy

Registering surfaces to the MNI template is, in principle, not necessary when quantifying the overlap of cortical representation using a relative measure (in our case, the intersect divided by the union of the corresponding surface areas). When comparing the respective centroids (cf. equation 1) or the centre-of-gravity (estimated via the points/locations of stimulations), standardising anatomy is mandatory, especially for between group comparisons. We realise that this procedure is not yet common in TMS-studies, whereas it is in, e.g., (f)MRI. For this reason we also conducted the same analyses on subject-specific MRIs without warping to the MNI-template. The corresponding results are reported as Supplementary Material S6 (in S7 we also show the case without constraining the analysis to M1). The results largely agree, at least in a qualitative form. This was clearly expected for our meaure of the relative overlap size (see above), but also for the centroids given our within-subject statistical design. The quantitative differences stem from the fact that the surface meshes of the subject-specific MRIs and the MNI template differ in resolution (triangularisation), which cannot be circumvented. Given this large agreement we are convinced that our approach to standardising antomy is proper.

### Limitations

While random coil positioning in combination with neuro-navigation is becoming common practise, the group analysis via transferring areas of excitability on a template is new. In fact, transforming centroid positions to the template is straightforward. Here, we used inflated spheres in combination with the great circle distance (see [28] for an alternative, namely, the minimisation Euclidian distances in deep brain regions). While this step is common in MRI studies, one should realise that its appropriateness stands and falls with the quality of surface inflation, and this might be difficult to quantify. Yet, we believe that this position is more accurate than (piecewise) affine transforms using isolated anatomical landmarks. However, when warping areas (and likewise volumes), geometrical transforms locally alter the neural density rendering subsequent biophysical modelling a challenge. As such, the ‘real’ degree of excitability should be interpreted with care and future studies should look in more detail into the possible effects. And, since the curvature between original and template surface may differ substantially, corrections for coil-orientation may become important.

A similar concern appears when recognising that the Intensity of TMS strongly influences the amount of overlap. Tardelli et al. [18] and DeJong et al. [22] already showed the relationship between the amount of overlap and intensity and noted that a higher stimulation intensity would result in a greater cortical overlap due to the non-focal characteristics of TMS. Especially at higher stimulation intensities, the observed overlap may – and probably will – be due to the stimulation at one point “radiating” to the adjacent surface, which will induce an MEP in other muscles. Any statement about the achieved spatial resolution when estimating areas (or volumes) of excitability and, more so, overlaps should therefore be questioned. To minimise this ‘radiating’ effect, we deliberately chose a minimum stimulation intensity just above RMT. We must admit that we cannot exclude radiation. Estimating the electrical field distributions [28-31] may help solving this, in particular when combined with other imaging modalities like fMRI [32]. The electrical field distribution could more informative than the focal point of the magnetic field (and its orientation). However, estimating requires several assumptions of the corresponding bio-electrical properties, first and foremost, that of grey matter, but we certainly advocate such multi-modal approaches for future studies.

## Conclusion

We used TMS to assess amount and position of the overlap between the cortical representations of eight hand and forearm muscles. We projected individual subject data to a high-resolution template cortical mesh. Hand muscles turned out to be more laterally positioned than the more medial forearm representations. Most synergistic muscle pairs displayed significantly more cortical overlap than their non-synergist counterparts.

## Supporting information

Supplementary Section

## Declaration of interest

The authors declare that they have no known competing financial interests or personal relationships that could have appeared to influence the work reported in this paper.

## Acknowledgements

FJ would like to thank the Chinese Scholarship Council for financial support (CSC # 201706210060). SMB was funded by a VIDI grant (no. 016.Vidi.178.014) from the Dutch Organization for Scientific Research (NWO).

### Appendix

**Figure A1.**
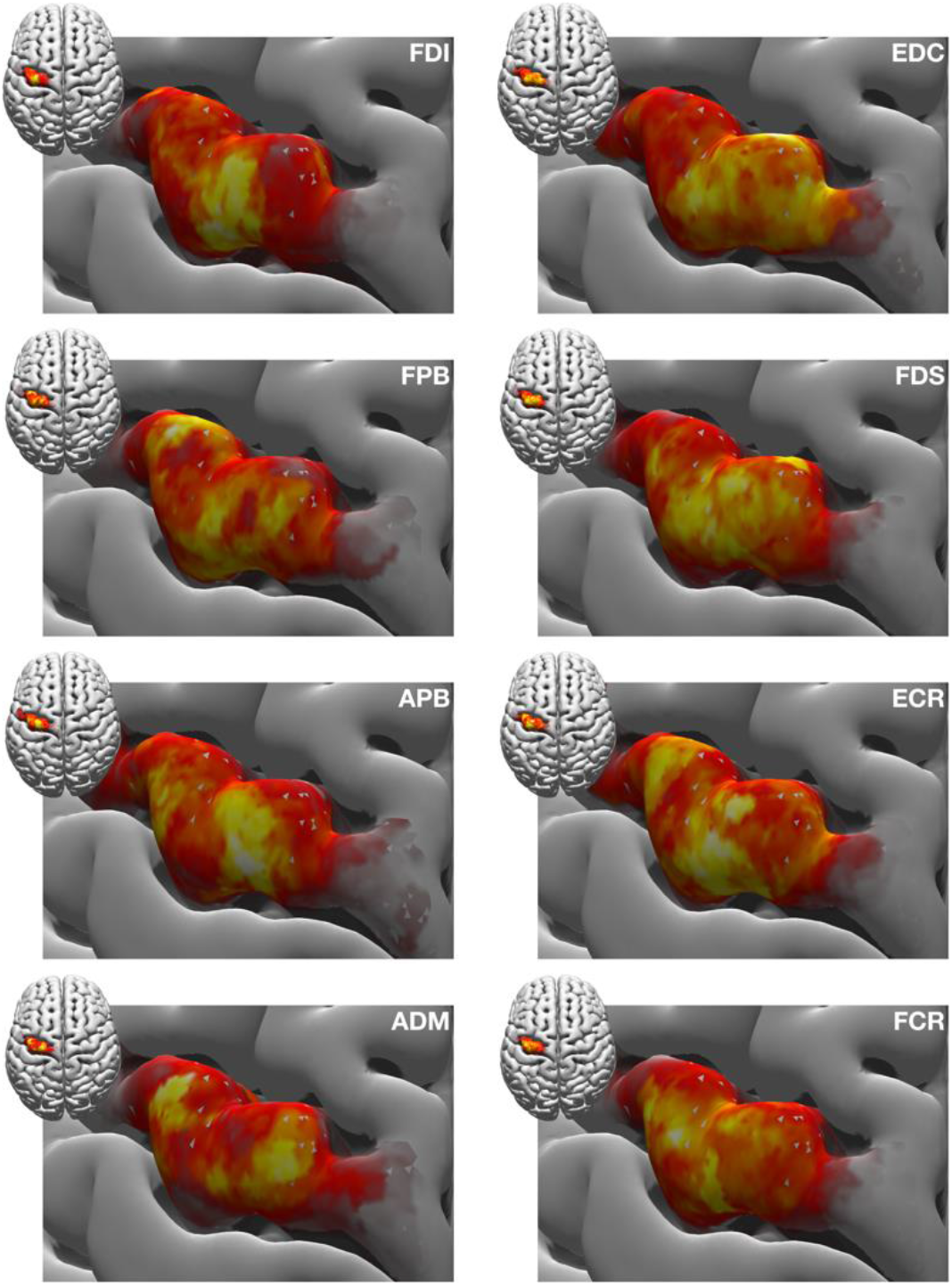
Group average of the cortical representations for the 105%-RMT-FDI Intensity. This figure agrees with Fig. 5 except for the colour-coding. While in Figure 5 intensities (MEP amplitudes) are given on a common scale, here every muscle map is scaled separately to highlight that the cortical distributions are typically complex even when looking at isolated muscles. Note that the distributions of excitability are not unimodal. There are in fact distinct peaks of excitability rendering the representation heterogeneous is not (solely) caused by averaging over the group. The units of the colour coding are arbitrary

## Notes

### Competing Interest Statement

The authors have declared no competing interest.

### Summary of Updates

revised according to comments from reviewers from Neuroimage: Reports

