## Supplementary Section for "Overlap in the cortical representation of hand and forearm muscles as assessed by navigated TMS"

### S1. Synergistic muscle pairs

**Table S1.1** Synergistic pairing based on the literature. Hand muscles are represented as light orange cells, and forearm muscles are represented by light blue ones. Dark (orange (hand-hand), green (hand-forearm), blue (forearm-forearm)) cells represent synergistic muscle pairs, the reference in the cell indicates which study suggested that that muscle pair is synergistic.

| | *FDI* | *ADM* | *APB* | *FPB* | *EDC* | *FDS* | *ECR* | *FCR* |
|---|---|---|---|---|---|---|---|---|
| *FDI* | | | [1,2] | [2] | [2] | [1,2] | | |
| *ADM* | | | | | [3] | [4] | | |
| *APB* | | | | [5] | [5] | [5] | | |
| *FPB* | | | | | | [5] | | |
| *EDC* | | | | | | | [6] | |
| *FDS* | | | | | | | | [7] |
| *ECR* | | | | | | | | |
| *FCR* | | | | | | | | |

### S2. Averaged cortical representations

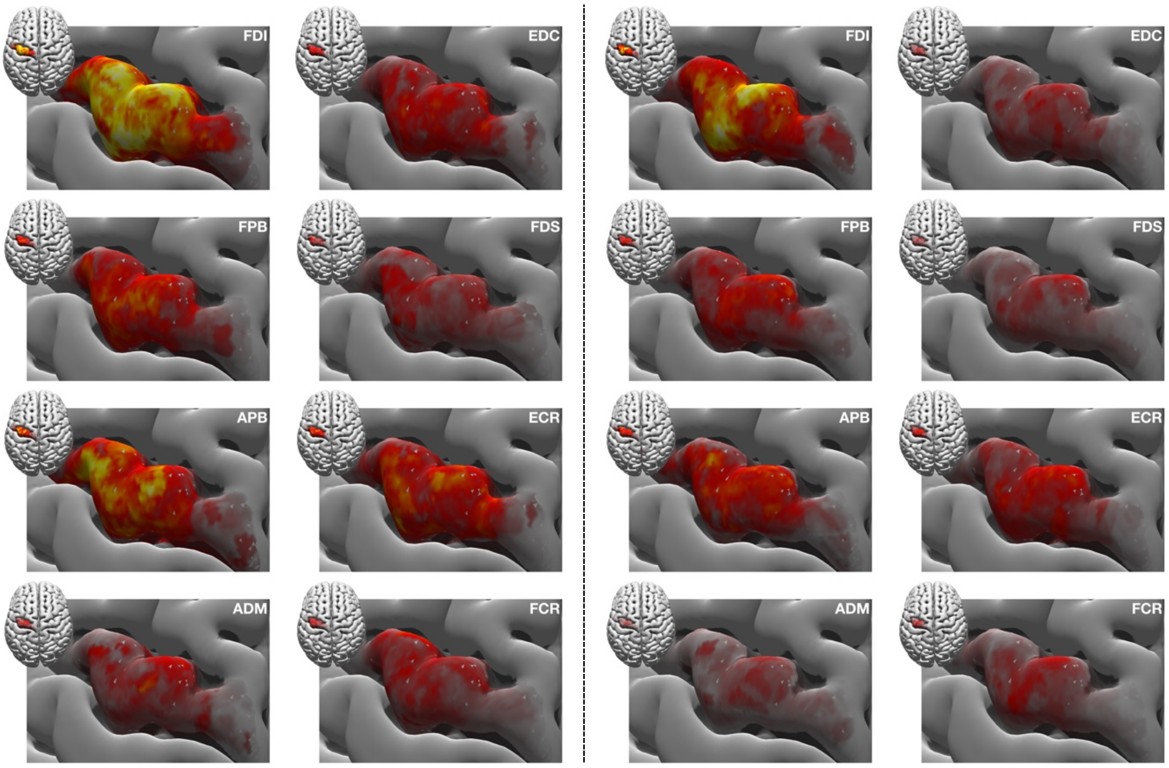


**Figure S2.1** Group average of the cortical representations for the 105%-RMT-EDC and the 105%-RMT-FCR intensities (left and right columns, respectively). As in Figure 5 of the main text also here the excitability of the FDI is most pronounced.

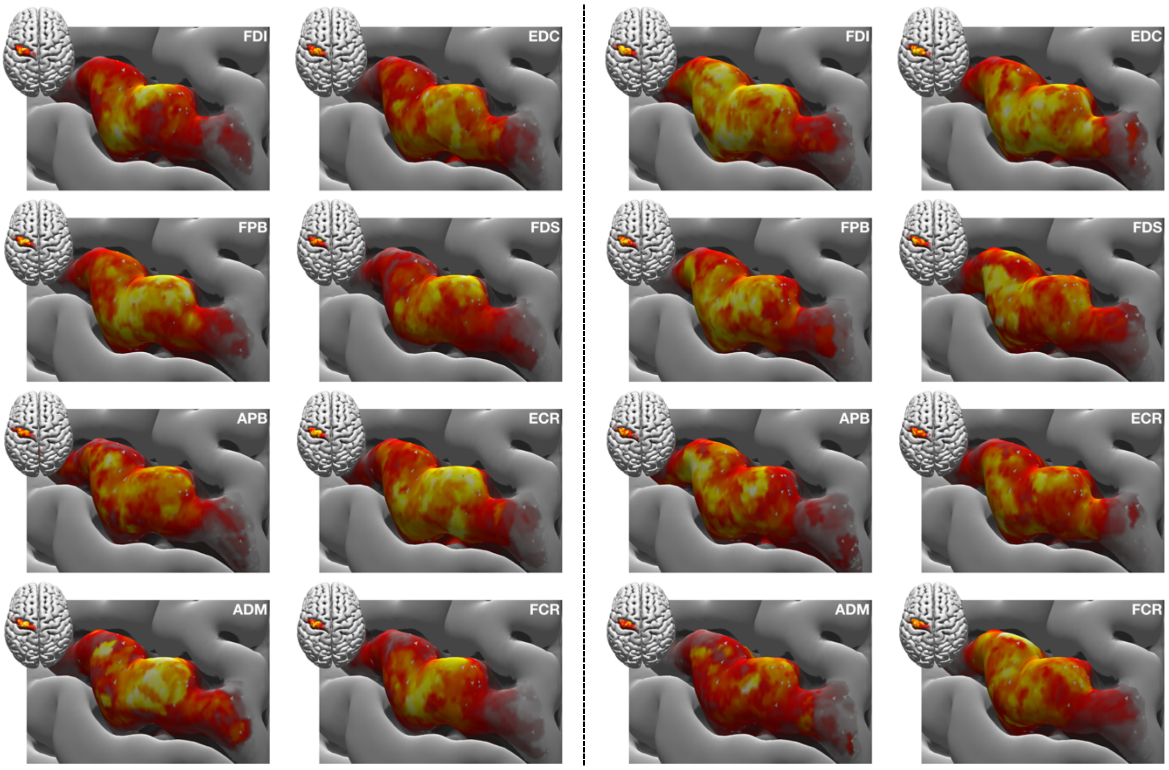


**Figure S2.2** Identical to Figure S2.1 but muscle intensities are display on individual colour scales; cf. Figure A1 in the main text.

### S3. Cortical representations of randomly selected subjects

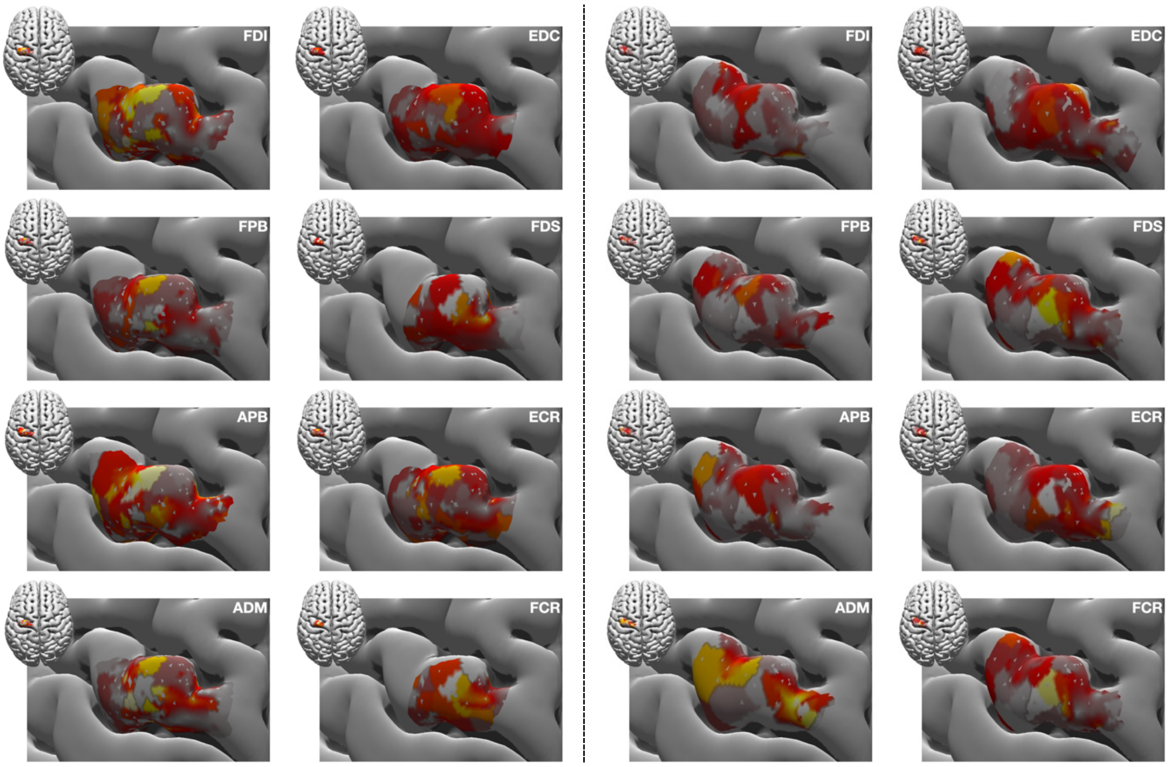


**Figure S3.1** Excitability maps at 105%-RMT-FDI of two randomly selected participants; obviously the between-subject variability is large despite the warping to the MNI template.

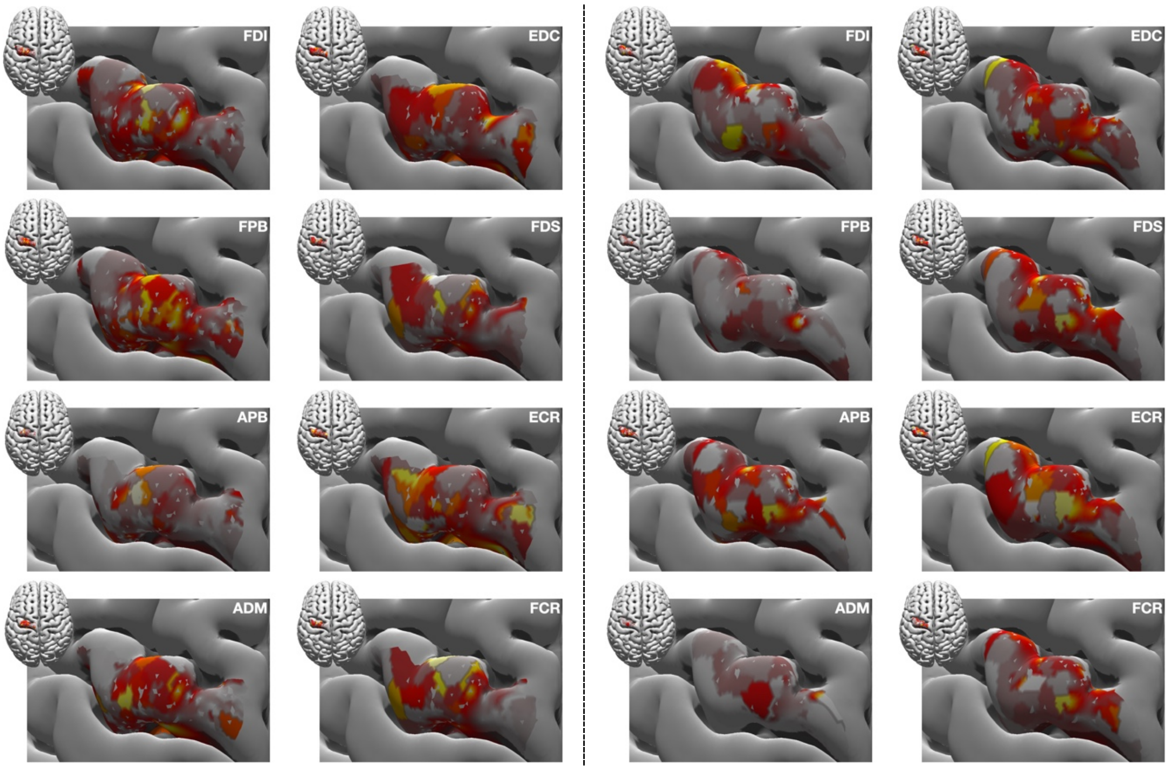


**Figure S3.2** Identical to Figure S3.1 but for the 105%-RMT-EDC intensity.

### S4. Centroid distributions in the posterior/anterior – lateral/medial plane

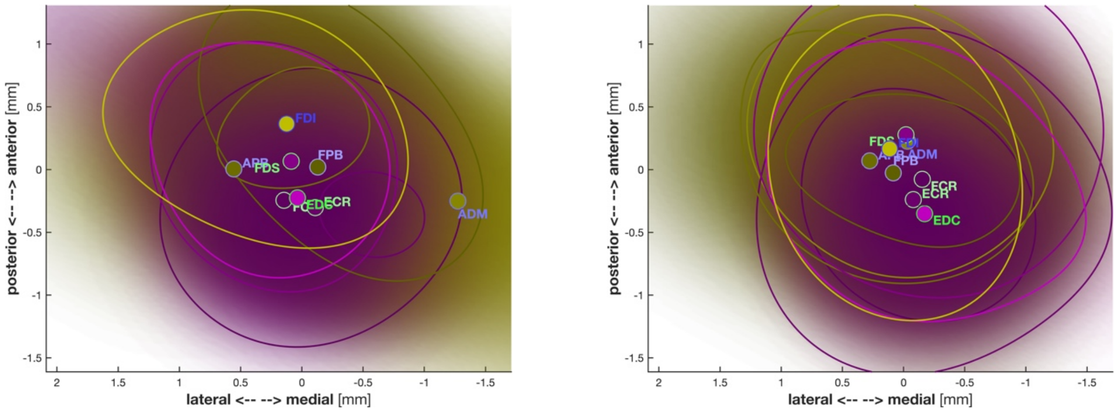


**Figure S4.1** Centroids averaged for all participants at 105%-RMT-EDC and -FCR intensities (left and right panel, respectively); cf. Figure 6 in the main text. Centroids clearly differ for different intensities rendering them less appropriate to parametrise muscle-specific areas; cf. main text, Figure 6.

### S5. Overlap statistics

**Table S5.1** Statistics of overlaps and centroids between the three groups of muscle pairs. $O_{kl}$ denotes the relative size of the pair-wise overlap and $C_{kl,x\ldots z}$ the corresponding centroids in medial/lateral, anterior/posterior, and inferior/superior directions, respectively. Significant effects are in **bold.**

| | Intensity | | Group | | Intensity × Group | |
|---|---|---|---|---|---|---|
| | F-value | *p*-value | F-value | *p*-value | F-value | *p*-value |
| $O_{kl}$ | F(2,38)=6.246 | **.005** | F(2,38)=208.862 | **.000** | F(4,76)=5.326 | **.001** |
| $C_{kl,x}$ | F(2,26)=0.366 | .697 | F(2,26)=156.069 | **.000** | F(4,52)=0.297 | .771 |
| $C_{kl,y}$ | F(2,26)=0.052 | .949 | F(2,26)=377.245 | **.000** | F(4,52)=0.281 | .790 |
| $C_{kl,z}$ | F(2,26)=0.874 | .429 | F(2,26)=12176.4 | **.000** | F(4,52)=0.972 | .390 |

**Table S5.2** Hand-hand pairs using the MNI template (constrained to M1). Significant effects in **bold**.

| | Intensity | | Pair | | Intensity × Pair | |
|---|---|---|---|---|---|---|
| | F-value | *p*-value | F-value | *p*-value | F-value | *p*-value |
| $O_{kl}$ | F(2,38)=4.979 | **.012** | F(5,95)=4.021 | **.034** | F(10,190)=0.422 | .819 |
| $C_{kl,x}$ | F(2,26)=0.594 | .559 | F(5,65)=0.759 | .468 | F(10,130)=1.081 | .377 |
| $C_{kl,y}$ | F(2,26)=0.282 | .756 | F(5,65)=4.023 | **.012** | F(10,130)=0.557 | .738 |
| $C_{kl,z}$ | F(2,26)=0.353 | .706 | F(5,65)=4.830 | **.012** | F(10,130)=0.530 | .696 |

**Table S5.3** Same as Table S5.2 but for the forearm-forearm muscle pairs.

| | Intensity | | Pair | | Intensity × Pair | |
|---|---|---|---|---|---|---|
| | F-value | *p*-value | F-value | *p*-value | F-value | *p*-value |
| $O_{kl}$ | F(2,38)=3.768 | **.032** | F(5,95)=3.870 | **.016** | F(10,190)=0.433 | .808 |
| $C_{kl,x}$ | F(2,26)=0.273 | .763 | F(5,65)=1.599 | .223 | F(10,130)=1.138 | .346 |
| $C_{kl,y}$ | F(2,26)=0.142 | .868 | F(5,65)=3.162 | **.043** | F(10,130)=0.771 | .503 |
| $C_{kl,z}$ | F(2,26)=1.400 | .265 | F(5,65)=0.494 | .670 | F(10,130)=0.897 | .467 |

**Table S5.4** Same as Tables S5.2&3 but for the hand-forearm muscle pairs.

| | Intensity | | Pair | | Intensity × Pair | |
|---|---|---|---|---|---|---|
| | F-value | *p*-value | F-value | *p*-value | F-value | *p*-value |
| $O_{kl}$ | F(2,38)=6.987 | **.003** | F(15,285)=1.859 | .164 | F(30,570)=0.668 | .706 |
| $C_{kl,x}$ | F(2,26)=0.342 | .714 | F(15,195)=1.489 | .229 | F(30,390)=1.150 | .343 |
| $C_{kl,y}$ | F(2,26)=0.045 | .956 | F(15,195)=4.214 | **.005** | F(30,390)=1.062 | .388 |
| $C_{kl,z}$ | F(2,26)=0.848 | .440 | F(15,195)=3.435 | **.012** | F(30,390)=0.624 | .672 |

### S6. Statistics using subject-specific MRIs constrained to M1

**Table S6** Statistics of the cortical muscle representation using subject-specific MRIs constrained to M1. $W$ = area size, centroid components: $C_x$ = medial/lateral, $C_y$ = anterior/posterior, $C_z$ = inferior/superior. The results agree qualitatively with the ones summarise in Table 1. Significant effects are in **bold**.

| | Intensity | | Muscle | | Intensity × Muscle | |
|---|---|---|---|---|---|---|
| | F-value | *p*-value | F-value | *p*-value | F-value | *p*-value |
| $C_x$ | F(2,32)=0.723 | .493 | F(7,112)=3.087 | **.021** | F(14,224)=1.369 | .235 |
| $C_y$ | F(2,32)=0.062 | .940 | F(7,112)=3.238 | **.004** | F(14,224)=0.725 | .748 |
| $C_z$ | F(2,32)=1.084 | .334 | F(7,112)=5.481 | **.000** | F(14,224)=0.893 | .506 |
| $W$ | F(2,30)=3.542 | .072 | F(7,105)=3.661 | **.029** | F(14,210)=1.481 | .246 |

**Table S6.1** Same as Table S5.1 but using the subject-specific MRI constrained to M1.

| | Intensity | | Group | | Intensity × Group | |
|---|---|---|---|---|---|---|
| | F-value | *p*-value | F-value | *p*-value | F-value | *p*-value |
| $O_{kl}$ | F(2,38)=5.543 | **.008** | F(2,38)=272.4530 | **.000** | F(4,76)=4.058 | **.013** |
| $C_{kl,x}$ | F(2,28)=0.573 | .570 | F(2,28)=106.0840 | **.000** | F(4,56)=1.461 | .226 |
| $C_{kl,y}$ | F(2,28)=0.308 | .738 | F(2,28)=447.8270 | **.000** | F(4,56)=0.407 | .675 |
| $C_{kl,z}$ | F(2,28)=0.648 | .470 | F(2,28)=4434.2650 | **.000** | F(4,56)=0.796 | .444 |

**Table S6.2** Same as Table S5.2 but using the subject-specific MRI constrained to M1.

| | Intensity | | Pair | | Intensity × Pair | |
|---|---|---|---|---|---|---|
| | F-value | *p*-value | F-value | *p*-value | F-value | *p*-value |
| $O_{kl}$ | F(2,38)=2.643 | .084 | F(5,95)=4.794 | **.021** | F(10,190)=0.507 | .746 |
| $C_{kl,x}$ | F(2,28)=1.557 | .229 | F(5,70)=1.851 | .168 | F(10,140)=0.298 | .851 |
| $C_{kl,y}$ | F(2,28)=0.839 | .443 | F(5,70)=1.091 | .358 | F(10,140)=0.701 | .552 |
| $C_{kl,z}$ | F(2,28)=0.241 | .695 | F(5,70)=5.027 | **.023** | F(10,140)=0.730 | .528 |

**Table S6.3** Same as Table S5.3 but using the subject-specific MRI constrained to M1.

| | Intensity | | Pair | | Intensity × Pair | |
|---|---|---|---|---|---|---|
| | F-value | *p*-value | F-value | *p*-value | F-value | *p*-value |
| $O_{kl}$ | F(2,38)=3.602 | **.037** | F(5,95)=5.774 | **.001** | F(10,190)=0.659 | .608 |
| $C_{kl,x}$ | F(2,28)=0.488 | .619 | F(5,70)=3.379 | **.035** | F(10,140)=0.335 | .789 |
| $C_{kl,y}$ | F(2,28)=0.057 | .945 | F(5,70)=4.064 | **.011** | F(10,140)=0.710 | .565 |
| $C_{kl,z}$ | F(2,28)=0.965 | .362 | F(5,70)=2.308 | .090 | F(10,140)=0.759 | .573 |

**Table S6.4** Same as Table S5.4 but using the subject-specific MRI constrained to M1.

| | Intensity | | Pair | | Intensity × Pair | |
|---|---|---|---|---|---|---|
| | F-value | *p*-value | F-value | *p*-value | F-value | *p*-value |
| $O_{kl}$ | F(2,38)=6.237 | **.005** | F(15,285)=1.63 | .199 | F(30,570)=0.638 | .719 |
| $C_{kl,x}$ | F(2,28)=0.623 | .543 | F(15,210)=2.642 | .053 | F(30,420)=0.446 | .844 |
| $C_{kl,y}$ | F(2,28)=0.31 | .736 | F(15,210)=2.078 | .103 | F(30,420)=0.854 | .513 |
| $C_{kl,z}$ | F(2,28)=0.674 | .463 | F(15,210)=3.902 | **.013** | F(30,420)=0.874 | .498 |

Tables S5.1…4 and S6.1…4 largely agree as expected given our statistical design (intensity, muscle, etc. are paired within subject). Qualitative differences are only incidental and occur solely in centroid components arguably due to difference in surface mesh resolution.

### S7. Statistics using subject-specific MRIs without constraining to M1

**Table S7** Same as Table S6 but without constraining the subject-specific MRIs to M1.

| | Intensity | | Muscle | | Intensity × Muscle | |
|---|---|---|---|---|---|---|
| | F | *p*-value | F | *p*-value | F | *p* |
| $C_x$ | F(2,34)=0.963 | .392 | F(7,119)=3.491 | **.008** | F(14,238)=0.573 | .726 |
| $C_y$ | F(2,34)=0.417 | .663 | F(7,119)=1.598 | .143 | F(14,238)=0.813 | .576 |
| $C_z$ | F(2,34)=1.644 | .208 | F(7,119)=5.853 | **.000** | F(14,238)=0.551 | .776 |
| $W$ | F(2,32)=7.237 | **.010** | F(7,112)=3.685 | **.036** | F(14,224)=2.522 | .080 |

**Table S7.1** Same as Table S6.1 but without constraining the subject-specific MRIs to M1.

| | Intensity | | Group | | Intensity × Group | |
|---|---|---|---|---|---|---|
| | F-value | *p*-value | F-value | *p*-value | F-value | *p*-value |
| $O_{kl}$ | F(2,38)=4.735 | **.015** | F(2,38)=388.614 | **.000** | F(4,76)=3.147 | **.043** |
| $C_{kl,x}$ | F(2,32)=1.475 | .244 | F(2,32)=106.019 | **.000** | F(4,64)=1.259 | .299 |
| $C_{kl,y}$ | F(2,32)=0.009 | .991 | F(2,32)=374.543 | **.000** | F(4,64)=0.441 | .702 |
| $C_{kl,z}$ | F(2,32)=0.686 | .511 | F(2,32)=4271.004 | **.000** | F(4,64)=0.969 | .407 |

**Table S7.2** Same as Table S6.2 but without constraining the subject-specific MRIs to M1.

| | Intensity | | Pair | | Intensity × Pair | |
|---|---|---|---|---|---|---|
| | F-value | *p*-value | F-value | *p*-value | F-value | *p*-value |
| $O_{kl}$ | F(2,38)=2.586 | .107 | F(5,95)=6.590 | **.008** | F(10,190)=0.414 | .840 |
| $C_{kl,x}$ | F(2,32)=0.780 | .467 | F(5,80)=1.694 | .195 | F(10,160)=0.760 | .564 |
| $C_{kl,y}$ | F(2,32)=0.431 | .654 | F(5,80)=2.580 | .069 | F(10,160)=0.210 | .929 |
| $C_{kl,z}$ | F(2,32)=0.593 | .559 | F(5,80)=8.773 | **.001** | F(10,160)=0.528 | .657 |

**Table S7.3** Same as Table S6.3 but without constraining the subject-specific MRIs to M1.

| | Intensity | | Pair | | Intensity × Pair | |
|---|---|---|---|---|---|---|
| | F-value | *p*-value | F-value | *p*-value | F-value | *p*-value |
| $O_{kl}$ | F(2,38)=4.055 | **.025** | F(5,95)=4.048 | **.010** | F(10,190)=0.956 | .440 |
| $C_{kl,x}$ | F(2,32)=1.493 | .240 | F(5,80)=1.289 | .289 | F(10,160)=0.874 | .470 |
| $C_{kl,y}$ | F(2,32)=0.123 | .885 | F(5,80)=0.411 | .655 | F(10,160)=0.870 | .486 |
| $C_{kl,z}$ | F(2,32)=0.811 | .453 | F(5,80)=2.157 | .096 | F(10,160)=0.666 | .550 |

**Table S7.4** Same as Table S6.4 but without constraining the subject-specific MRIs to M1.

| | Intensity | | Pair | | Intensity × Pair | |
|---|---|---|---|---|---|---|
| | F-value | *p*-value | F-value | *p*-value | F-value | *p*-value |
| $O_{kl}$ | F(2,38)=4.707 | **.015** | F(15,285)=2.197 | .112 | F(30,570)=1.047 | .404 |
| $C_{kl,x}$ | F(2,32)=1.602 | .217 | F(15,240)=1.651 | .176 | F(30,480)=0.541 | .790 |
| $C_{kl,y}$ | F(2,32)=0.004 | .996 | F(15,240)=1.379 | .252 | F(30,480)=0.519 | .768 |
| $C_{kl,z}$ | F(2,32)=0.758 | .477 | F(15,240)=4.510 | **.004** | F(30,480)=0.553 | .739 |

As expected, constraining stimulations to M1 has a major effect on the statistics across results both for the overlaps and the centroid positions.

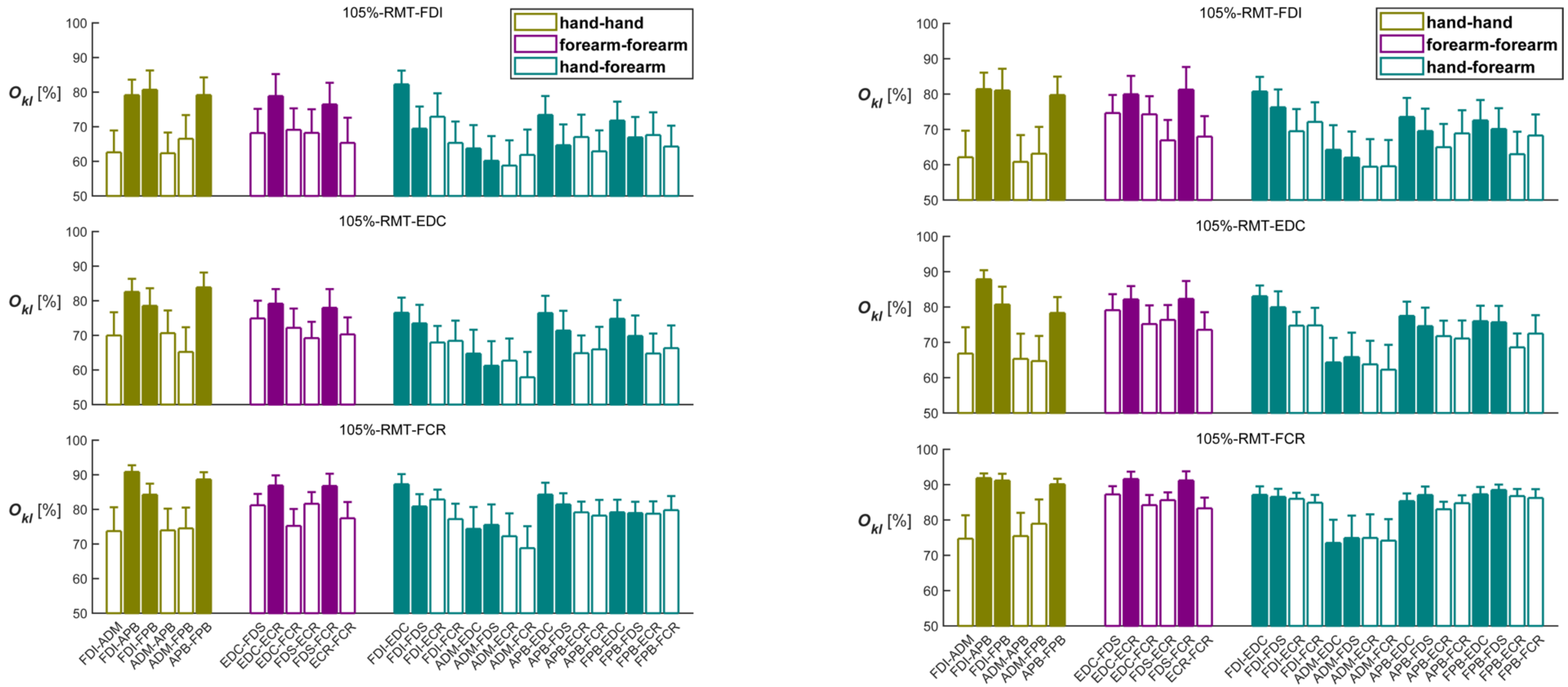


**Figure S7.1** Overlap of all muscle pairs using subject-specific MRIs (left column: constrained to M1; right column: without constraint). We show the relative overlaps $O_{kl}$ between muscle pairs expressed in percentage (of the union area) for the three intensities. The filled bars represent the muscle pairs with synergistic function, and the open bars represent the non-synergistic muscle pairs. Error bars represent the standard errors over participants. Both estimates agree qualitatively with the ones shown in Figure 7 of the main text, especially, when constraining to M1. This is expected due to the way we normalised $O_{kl}$. The minor quantitative differences stem from differences in resolution (triangularisation) of the cortex surfaces.
